# Metabolomics and biochemical approaches link salicylic acid biosynthesis to cyanogenesis in peach plants

**DOI:** 10.1101/147819

**Authors:** Diaz-Vivancos Pedro, Bernal-Vicente Agustina, Cantabella Daniel, Petri Cesar, Hernández José Antonio

**Author notes:** Corresponding author Dr. Pedro Diaz-Vivancos, CEBAS-CSIC, Campus Universitario de Espinardo, 25. 30100 Murcia (Spain).

## Abstract

**Highlight:** Mandelonitrile, and hence cyanogenic glycosides turnover, is involved in salicylic acid (SA) biosynthesis in peach plants under control and stress conditions. A third pathway for SA synthesis in peach is proposed.

**Abstract:** Despite the long-established importance of salicylic acid (SA) in plant stress responses and other biological processes, its biosynthetic pathway has not been fully characterized. The proposed SA synthesis originates from chorismate by two distinct pathways: isochorismate and penhylalanine (Phe) ammonia-lyase (PAL) pathways. Cyanogenesis is the process related to the release of hydrogen cyanide from endogenous cyanogenic glycosides (CNglcs), and it has been linked to plant plasticity improvement. To date, however, no relationship has been suggested between both pathways. In this work, by metabolomics and biochemical approaches (including [^13^C]-labelled compounds), we provide evidences showing that CNglcs turnover is involved, at least in part, in SA biosynthesis in peach plants under control and stress conditions.

The main CNglcs in peach are prunasin and amygdalin, with mandelonitrile (MD), synthesized from Phe, controlling their turnover. In peach plants MD is at the hub of the suggested new SA biosynthetic pathway and CNglcs turnover, regulating both the amygdalin and SA biosynthesis. MD-treated peach plants displayed increased SA levels via benzoic acid (SA precursor). In addition, MD also provides partial protection against *Plum pox virus* infection in peach seedlings. Thus, we proposed a third pathway, alternative to the PAL pathway, for SA synthesis in peach plants.

## Introduction

The plant hormone salicylic acid (SA) is the focus of intensive research due to its function as an endogenous signal mediating plant defense responses against both biotic and abiotic stimuli. In addition to its well-known role as a key signaling and regulatory molecule in plant defense responses, SA also plays crucial roles in diverse biological processes such as cell growth and development, seed germination, stomatal aperture, and fruit yield, among others (Liu *et al.*, 2015; Rivas-San Vicente and Plasencia, 2011). Most of the currently available information about the SA biosynthesis pathway comes from works on Arabidopsis and other herbaceous plants (Chen *et al.*, 2009; Dempsey *et al.*, 2011). The proposed SA synthesis originates from chorismate, the end product of the shikimate pathway, by two distinct pathways: the isochorismate (IC) and the penhylalanine (Phe) ammonia-lyase (PAL) pathways (Dempsey *et al.*, 2011). The PAL pathway uses Phe as substrate, but its contribution to the total SA pool is minimal (ca. 5% of the total SA synthesis). The IC pathway, on the other hand, accounts for the bulk of SA synthesis (ca. 95% of the total SA synthesis) (Chen *et al.*, 2009). Nevertheless, the biosynthetic pathway of SA in plants has not been fully characterized yet (Dempsey *et al.*, 2011), and knowledge regarding this topic is even scarcer in woody plants such as fruit trees.

Cyanogenic glycosides (CNglcs) are specialized plant compounds (derived from amino acids) that release toxic hydrogen cyanide (HCN) and ketones when hydrolyzed by β-glycosidases and α-hydroxynitrilases in a process referred to as cyanogenesis (Gleadow and Møller, 2014). The main cyanogenic glucosides in *Prunus* species are prunasin and amygdalin, with mandelonitrile (MD) at the hub of its turnover (Sánchez-Pérez *et al.*, 2008). Whereas CNglcs have traditionally been associated with protection against herbivore and fungal attack, their role in other biological processes such as germination and bud burst has been suggested (Gleadow and Møller, 2014). Nevertheless, endogenous CNglcs turnover may be highly species-dependent, and new functions for these molecules remain to be elucidated. For example, secondary metabolites turnover could act dissipating excess energy and providing reducing power in stress conditions (Neilson *et al.*, 2013; Selmar and Kleinwachter, 2013). Moreover, CNglcs may also be able to quench reactive oxygen species such as H_2_O_2_, suggesting a possible role for these glycosides during unfavorable environmental conditions (Gleadow and Møller, 2014).

In peach plants, MD synthesized from Phe via cytochrome P450 enzymes (CYP79 and CYP71) is converted into benzaldehyde and HCN by mandelonitrile lyase (MDL) activity, and benzaldehyde can be easily oxidized to produce benzoic acid (BA). In addition, benzaldehyde and benzoic acid appear as intermediate precursors of SA biosynthesis via the PAL pathway (Dempsey *et al.*, 2011; Ribnicky *et al.*, 1998). This fact led us to consider a relationship between cyanogenesis and SA biosynthesis (Fig. 1). Moreover, both SA and HCN are involved in thermogenesis events by the induction of the alternative oxidase pathway, either directly (SA) or through the inhibition of cytochrome c oxidase (HCN) (Taiz and Zeiger, 2010). In the present work, by feeding GF305 peach (*Prunus persica* L.) micropropagated shoots and seedlings with MD and Phe, we have accumulated strong evidence suggesting that MD could be metabolized into SA, linking the SA biosynthetic and cyanogenic glucoside pathways in peach plants. Here we show that MD could act as a hub controlling CNglcs turnover and, at least in part, SA biosynthesis. It therefore seems that a third pathway for SA synthesis is present in peach plants, being this pathway functional under both control and stress conditions. This pathway is an alternative to the PAL pathway for SA biosynthesis from Phe, and it is initiated by cytochrome P450 enzymes, similar to the indole-3-acetaldoxime pathway for auxin biosynthesis from tryptophan (Mano and Nemoto, 2012).

**Figure 1.**
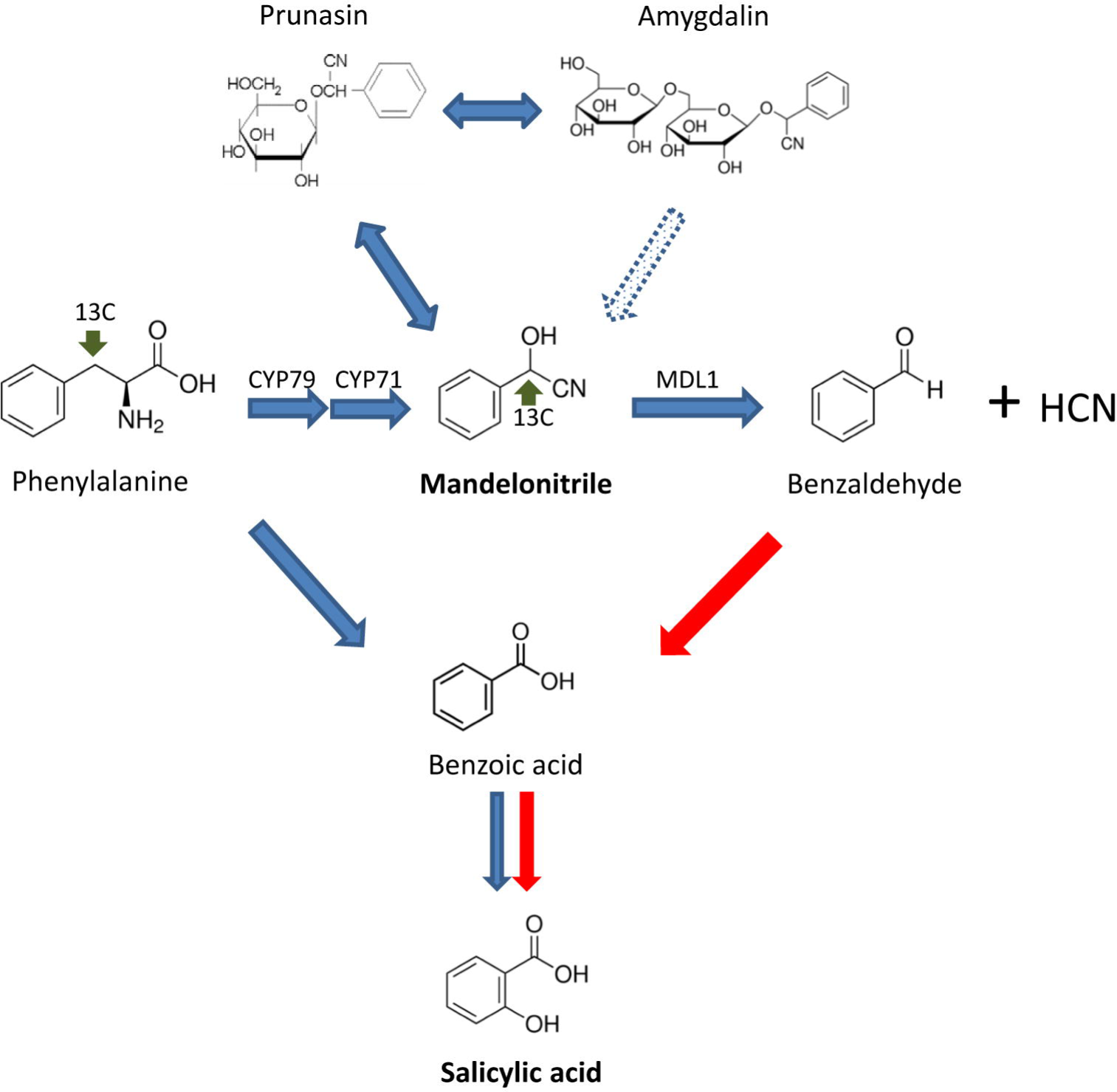
Proposed SA biosynthetic pathway in peach plants. Blue arrows indicate the already described SA biosynthesis in plants (dot arrow, putative; CYP79 and CYP71, Cyt P450 monooxygenases; MDL1, mandelonitrile lyase), whereas red arrows show the new pathway suggested for peach plants.

Furthermore, MD levels could be involved in defense responses by increasing the levels of SA and/or its interaction with oxidative signaling defense-induced pathways. To assess this hypothesis we have also analyzed the levels of enzymatic and non-enzymatic antioxidants in MD- and Phe-treated peach plants in addition to the expression of two genes involved in redox signaling and the phenotypic scoring system used for evaluating resistance and susceptibility to *Plum pox virus* (PPV) infection (Decroocq *et al.*, 2005).

## Material and Methods

### Plant material

The assays were performed on GF305 peach (*Prunus persica* L.) plants, both under greenhouse and *in vitro* conditions. For *ex vitro* assays, after submitting the GF305 peach seedlings to an artificial rest period in a cold chamber to ensure uniformity and fast growth, seedlings were grown in 2 L pots in an insect-proof greenhouse and distributed to 3 batches (control and MD- and Phe-treated) of 15 plants each. Two different experiments were carried out in 2015 and 2016. Due to the fact that in the soil a small proportion of MD could be dissociate non-enzymatically during the course of the experiment, plants were irrigated twice per week with water (control) and 1 mM MD or 1 mM Phe for 7 weeks.

For *in vitro* assays [^13^C]-labelled compounds were used, and 200 μM MD- or Phe-alpha[^13^C] (Campro Scientific GmbH, Germany) were added to the micropropagation media during two sub-cultures. The micropropagated GF305 peach shoots were sub-cultured at 4-week intervals for micropropagation (Clemente-Moreno *et al.*, 2011).

### Metabolomics analysis

The levels of Phe, MD, amygdalin, benzoic acid and SA were determined in *in vitro* micropropagated shoots at the Metabolomics Platform at CEBAS-CSIC (Murcia, Spain). Leaf samples from micropropagated shoots were extracted in 50% methanol, filtered in PTFE 0.45 μm filters (Agilent Technologies) and analyzed using an Agilent 1290 Infinity UPLC system coupled to a 6550 Accurate-Mass quadrupole TOF mass spectrometer (Agilent Technologies). Standard curves for each compound were performed, and data were processed using Mass Hunter Qualitative Analysis software (version B.06.00 Agilent Technologies). Hormone levels in the leaves of MD-treated GF305 seedlings were determined using a UHPLC-mass spectrometer (Q-Exactive, ThermoFisher Scientific) at the Plant Hormone Quantification Platform at IBMCP (Valencia, Spain).

### Extraction and enzymatic assays

*In vitro* shoots and *ex vitro* leaf samples were homogenized with an extraction medium (1:3, w/v) containing 50 mM Tris-acetate buffer (pH 6), 0.1 mM EDTA, 2 mM cysteine, 0.2% (v/v) Triton X-100, 2% (w/v) polyvinylpolypyrrolidone (PVPP) and 1% (w/v) polyviny-pyrrolidone (PVP). To determine APX, 20 mM ASC was added to the extraction medium. The extracts were filtered through two layers of nylon cloth and centrifuged at 13000 rpm for 10 min. The supernatant fraction was desalted on Sephadex G-25 NAP columns equilibrated with the same buffer used for homogenization. For APX activity, 2 mM sodium ascorbate was added to the equilibration buffer. The activities of the ASC–GSH cycle enzymes, POX, CAT and SOD, were assayed as previously described (Diaz-Vivancos *et al.*, 2008; Diaz-Vivancos *et al.*, 2013; Diaz-Vivancos *et al.*, 2006). MDL activity was assayed by monitoring the increase of absorbance at 280 nm due to the benzaldehyde released by the enzymatic hydrolysis of DL-mandelonitrile (Ueatrongchit *et al.*, 2008; Willeman *et al.*, 2000). The enzyme solution was added to a 50 mM Tris-acetate buffer pH 5 containing 0.1 mM mandelonitrile in a total volume of 1 ml. Protein determination was performed according to the method of Bradford (Bradford, 1976).

### Ascorbate and Glutathione analysis

Leaf samples were snap-frozen in liquid nitrogen and stored at -80°C until use. The frozen samples were homogenized in 1 ml 1 M HClO4. Homogenates were centrifuged at 12 000 g for 10 min, and the supernatant was neutralized with 5 M K2CO3 to pH 5.5-6. The homogenate was centrifuged at 12 000 g for 1 min to remove KClO4. The supernatant obtained was used for ascorbate and glutathione determination (Pellny *et al.*, 2009; Vivancos *et al.*, 2010).

### Gene expression

RNA samples were extracted using a GF1-Total RNA Extraction Kit (Vivantis) according to the manufacturer’s instructions. The expression levels of *MDL, NPR1*, *TrxH* and the reference gene *translation elongation factor II* (*TEF2*) (Tong *et al.*, 2009) were determined by real-time RT-PCR using the GeneAmp 7500 sequence detection system (Applied Biosystems, Foster City, CA, USA) (Faize *et al.*, 2013). The accessions and primer sequences are as follows: *MDL1* (Y08211.1; forward 5’- gtttcgcttgcaaagagggg-’3; reverse 5’-gctttagggagtcatttccttgc-’3); NPR1 (DQ149935; forward 5’-tgcacgagctcctttagtca’3; reverse 5’-cggcttactgcgatcctaag-’3); *TrxH* (AF323593.1; forward 5’- tggcggagttggctaagaag-’3; 5’-ttcttggcacccacaacctt-’3); and *TEF2* (TC3544; forward 5’- 0-’3; reverse 5’-gaaggagagggaaggtgaaag-’3). Relative quantification of gene expression was calculated by the Delta-Delta Ct method, and the expressions of the genes of interest were normalized with the endogenous control *TEF2*.

### Statistical analysis

The data were analyzed by one-way or two-way ANOVA using SPSS 22 software. Means were separated with the Duncan’s Multiple Range Test (P < 0.05). F-values and probabilities associated with the main effects and possible interactions are indicated where appropriate.

## Results

### SA biosynthesis from CNglcs

GF305 peach micropropagated shoots were fed with [^13^C]Phe or with [^13^C]MD. The addition of these compounds to the culture media had no important effect in the growth of the micropropagated peach shoots (Fig 2A). We determined the percentage of [^13^C]- labelled compounds from the total content of Phe, MD, BA and SA in the micropropagated peach shoots treated with either [^13^C]Phe or with [^13^C]MD (Fig. 3). Due to the high sensitivity of the UPLC-Quadrupole-TOF-MS system used for metabolomics analysis, we detected basal levels (less than 10%) of [^13^C]Phe, [^13^C]MD and [^13^C]SA in control micropropagated shoots (Fig. 3). It is important to note that we only found [^13^C]BA (one SA precursor) in non-stressed [^13^C]MD-fed micropropagated shoots. Of the total BA detected in [^13^C]MD-treated micropropagated shoots, nearly 40% was [^13^C]-labelled. In the presence of [^13^C]MD, nearly 20% of the total SA quantified appeared as [^13^C]SA. Regarding the [^13^C]Phe treatment, only a slight increase in the percentage of [^13^C]MD was observed in non-stressed *in vitro* peach shoots (Fig. 3).

**Figure 2.**
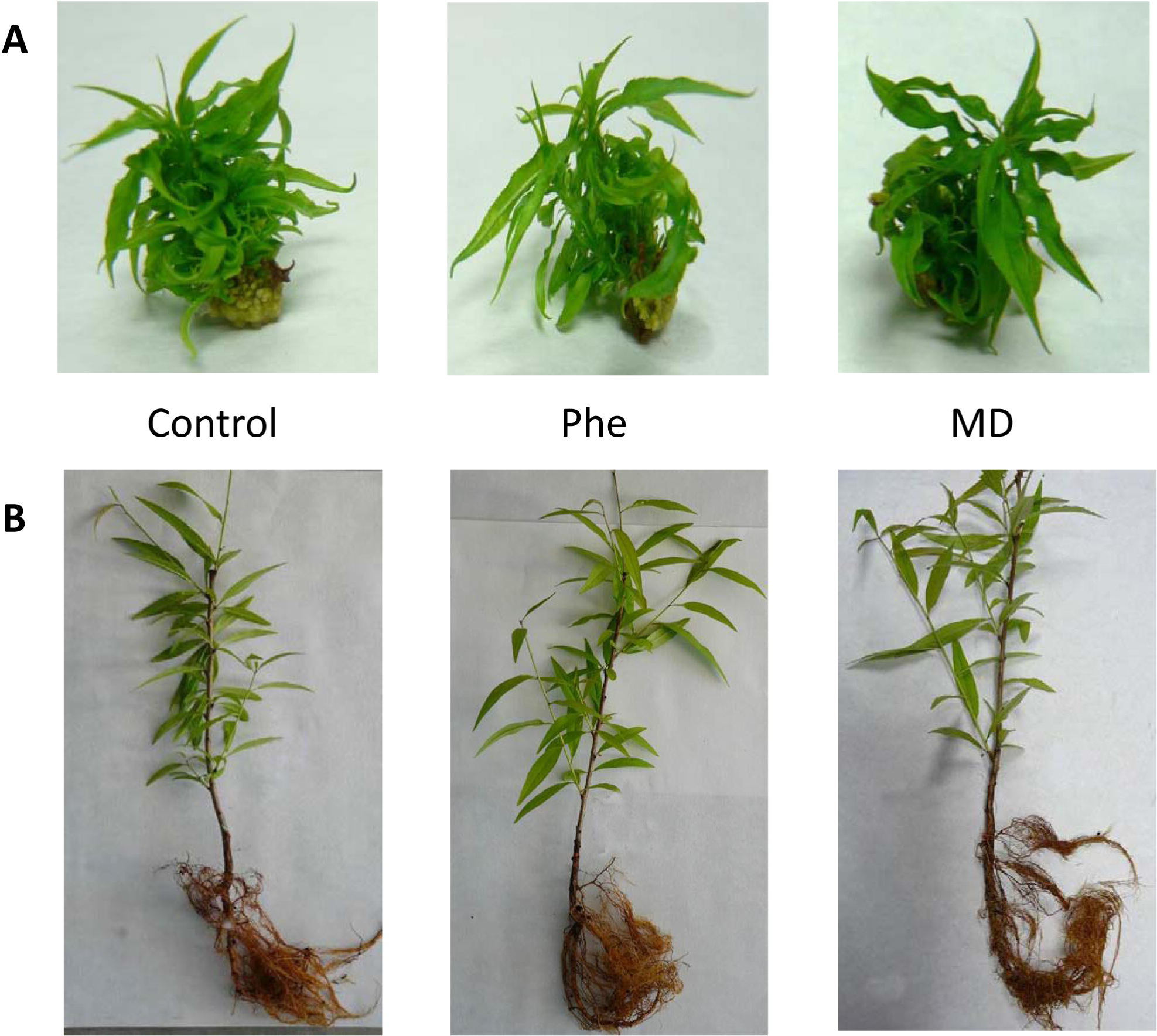
Micropropagated GF305 peach shoots (A) and seedlings (B) grown under control conditions and in the presence of mandelonitrile (MD) and phenylalanine (Phe). *In vitro* shots were micropropagated by incorporating 200 μM of either [^13^C]MD or [^13^C]Phe into the media, whereas peach seedlings were watered with 1 mM solution of MD or Phe. Neither of the treatments had effects on plant growth and development.

**Figure 3.**
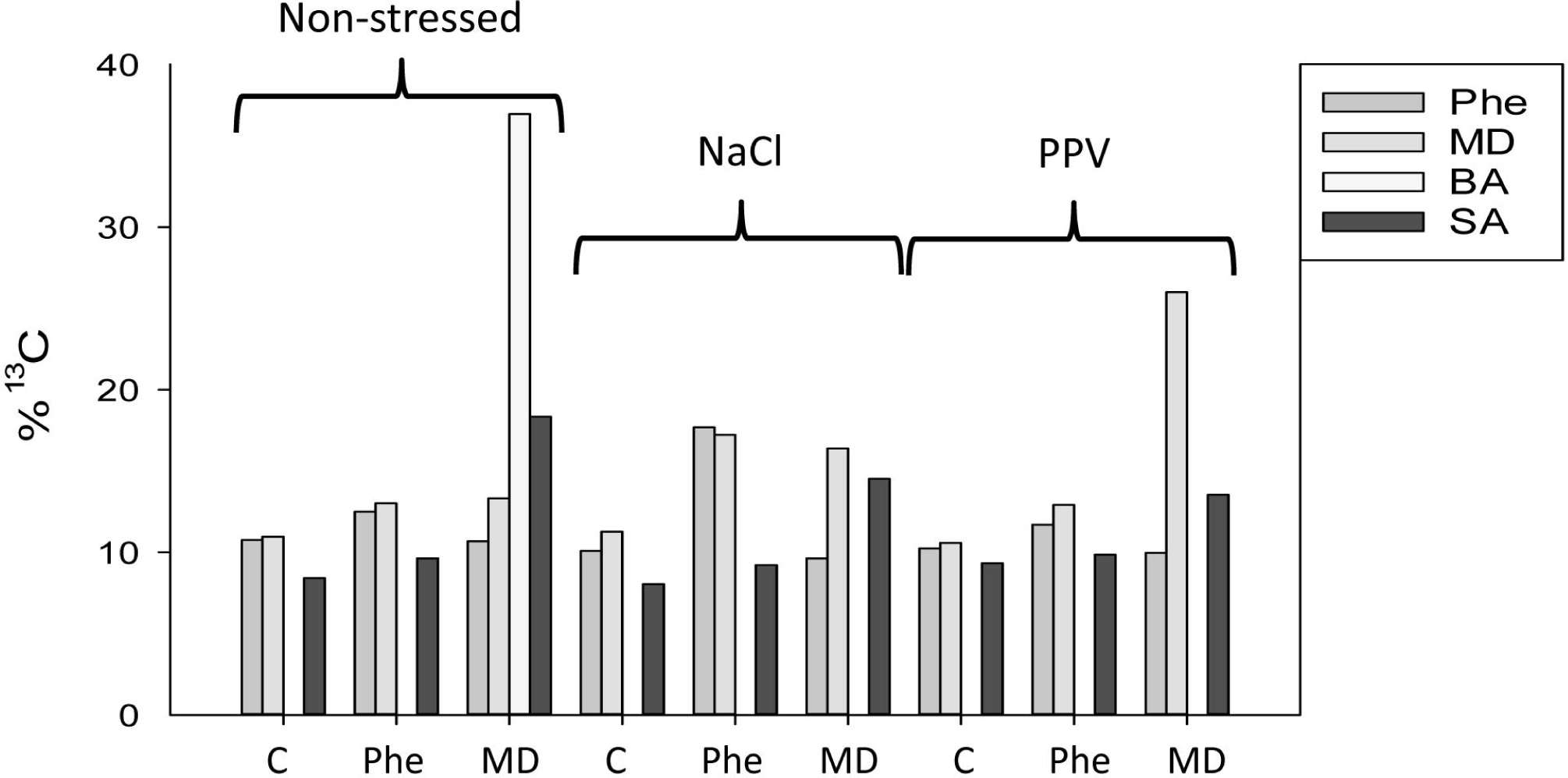
Percentage of [^13^C]-phenylalanine, mandelonitrile, benzoic acid, and salicylic acid in non-stressed, NaCl-stressed and PPV-infected peach shoots micropropagated in the presence or absence of [^13^C]MD or [^13^C]Phe. Under control conditions, basal levels of [^13^C]- mandelonitrile, phenylalanine and salicylic acid were observed, whereas [^13^C]benzoic acid was only detected in [13C]MD-treated micropropagated shoots. Ions with an additional 1.0035 accurate mass and confirmation by isotopic distribution and spacing were defined as ions marked with ^13^C. Data represent the mean of at least 20 repetitions of each treatment.

After treatment, [^13^C]Phe-fed micropropagated shoots showed a significant increase in amygdalin (61%) and a non-significant increase in BA (Fig. 4). The [^13^C]MD-fed micropropagated shoots displayed a similar increase in amygdalin to that produced by the [^13^C]Phe treatment, indicating that the CNglcs pathway is fully functional under our experimental conditions. Interestingly, however, significant increases in BA and SA (of about 80%) were only observed after the [^13^C]MD treatment (Fig. 4).

**Figure 4.**
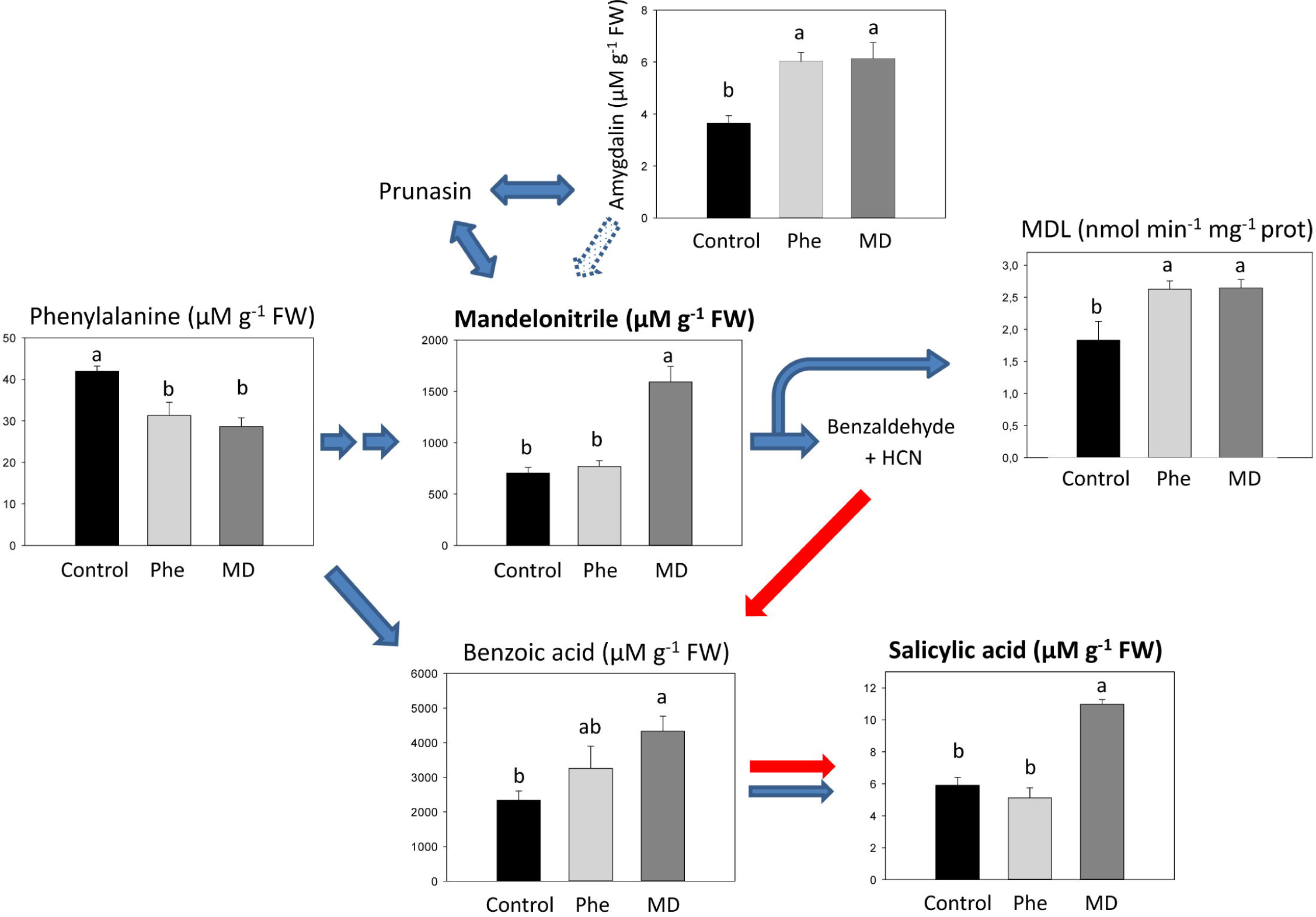
Total levels (μM g^-1^ FW) of amygdalin, benzoic acid, mandelonitrile, phenylalanine and salicylic acid, and MDL enzymatic activity in micropropagated peach shoots in the presence or absence of [^13^C]MD or [^13^C]Phe. Data represent the mean ± SE of at least 12 repetitions of each treatment. Different letters indicate significant differences in each graph according to Duncan’s test (P<0.05).

Given the effect of MD treatments on SA levels in micropropagated peach shoots, we also fed peach seedlings grown under normal physiological conditions in a greenhouse with either MD or Phe. As observed in micropropagated peach shoots, irrigation with MD or Phe had no significant effects on the growth of peach seedlings, which showed normal growth in both the shoots and roots (Fig 2B). In this experiment, we analyzed the effect of the 1 mM MD treatment on the hormone profile of leaf samples. Again, the MD treatment produced an increase in SA levels (about 88%), similar to that noticed in *in vitro* micropropagated peach shoots (Table 1). Furthermore, due to the well-known cross-talk among plant hormones, MD affected also the levels of other hormones. The amount of ABA, another stress-related plant hormone, increased up to 45%. In addition, the 1 mM MD treatment also produced a significant increase in both the gibberellin GA1 and the cytokinin dihidrozeatine (DHZ) in the same range (nearly 60%) (Table 1).

**Table 1.**
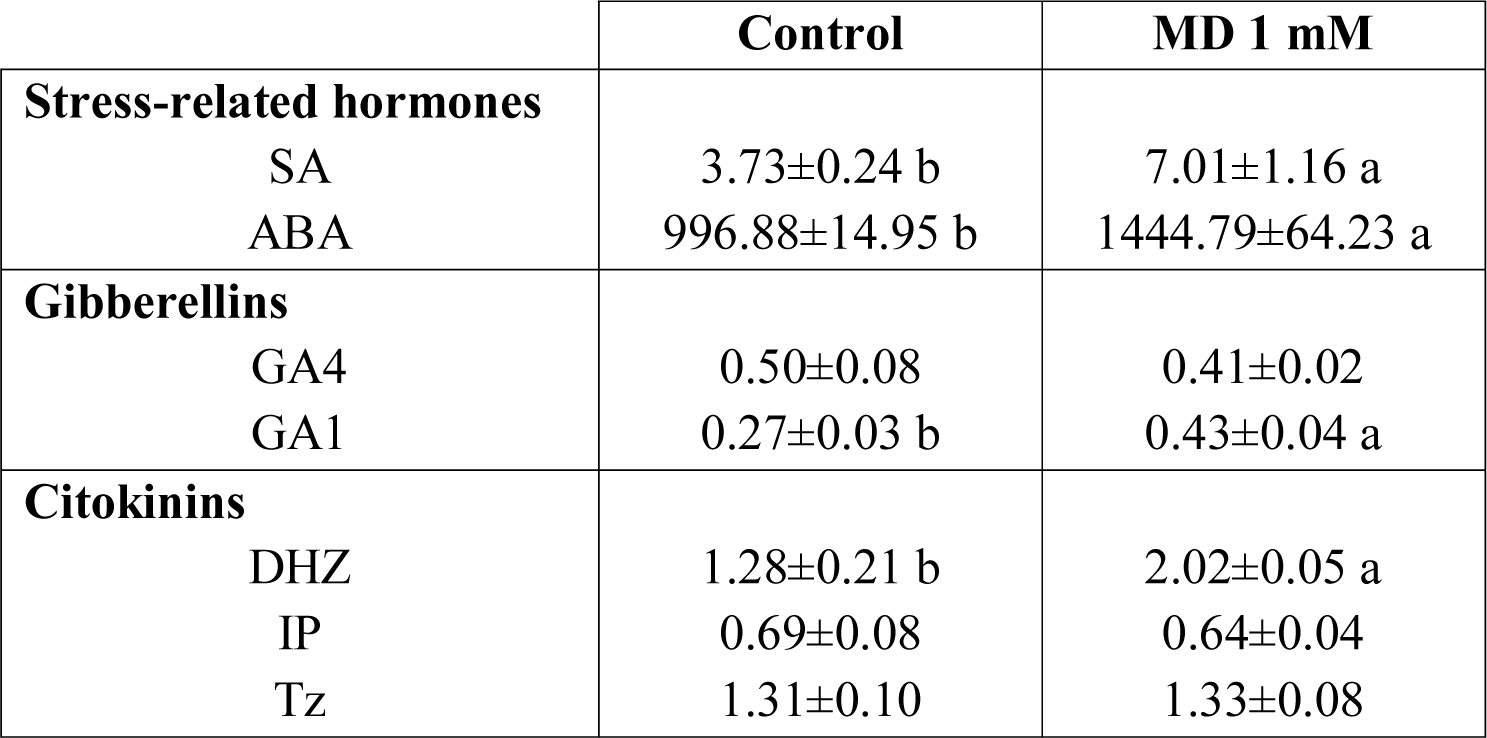
Effect of MD treatment on hormone levels in leaves of GF305 seedlings grown under greenhouse conditions. Data are expressed as ng g^-1^ FW except for SA levels (μg g^-1^ FW). Data represent the mean ± SE of at least eight repetitions of each treatment. Different letters indicate significant differences according to Duncan’s test (P<0.05).

### SA biosynthesis and plant performance under stress conditions

The level of SA was also determined in micropropagated peach shoots submitted to both abiotic and biotic stresses. Abiotic stress was achieved adding 30 mM NaCl to the culture media whereas *Plum pox virus* (PPV)-infected *in vitro* shoots (Clemente-Moreno *et al.*, 2011) were used to assess the biotic stress condition. PPV is the causal agent of sharka disease and the most destructive and detrimental disease affecting Prunus species (Clemente-Moreno *et al.*, 2015). Regarding SA biosynthesis, under control conditions a similar strong increase in the total content of SA was observed in NaCl- and PPV-stressed micropropagated peach shoots (Fig. 5). Contrary to what observed in non-stressed peach shoots, under both stress conditions [^13^C]Phe increased SA, whereas [^13^C]MD treatment did not increase the total SA content (Fig. 5). Nevertheless, stressed [^13^C]MD-fed micropropagated shoots displayed increases in the percentages of [^13^C]MD (45% and 148% by NaCl and PPV infection respectively) and [^13^C]SA (75% and 45% by NaCl and PPV infection respectively) (Fig. 3).

**Figure 5.**
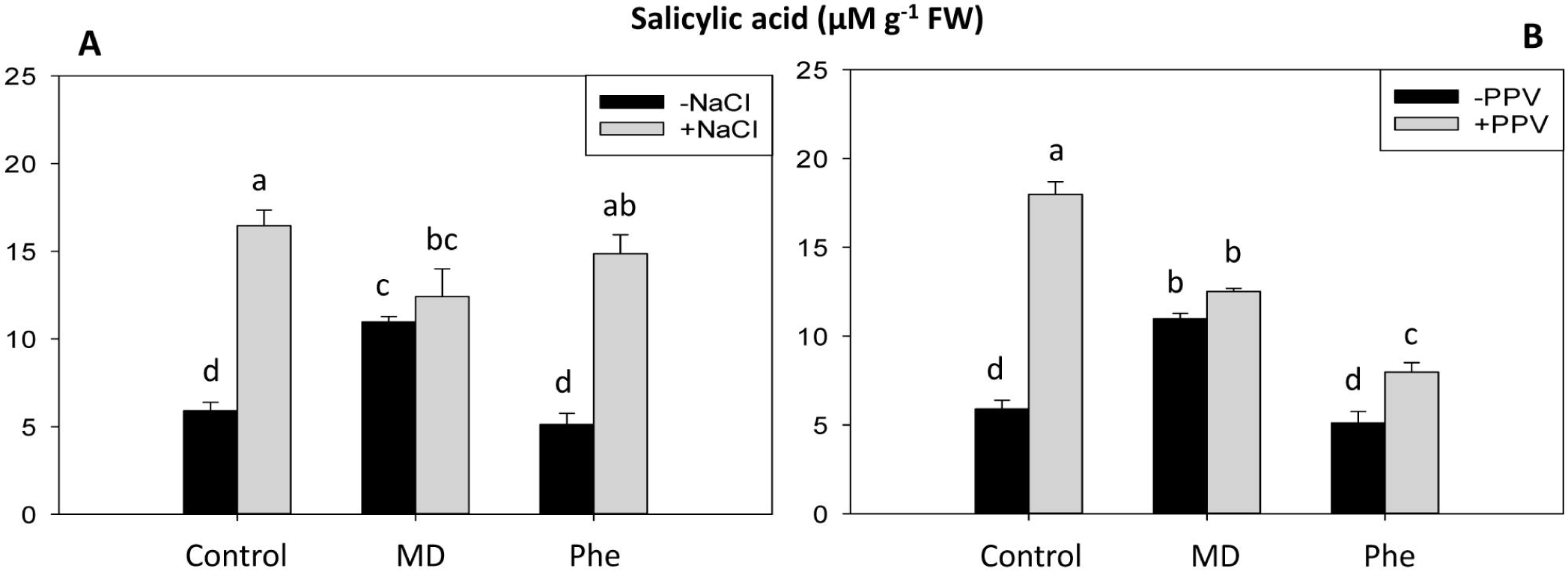
Total level (μM g^-1^ FW) of salicylic acid in peach shoots micropropagated in the presence or absence of [13C]MD or [13C]Phe, submitted to salt stress (30 mM NaCl; A) or PPV infection (B).

In addition, according to salinity damage observed, MD-treatment had not a significant effect on the *in vitro* shoots performance, whereas Phe seems to increase the salt stress deleterious effect, as observed by the increase of leaves showing salinity injure per micropropagated shoot (Fig. S1). On the other hand, it is well known that SA is a key signaling molecule involved in systemic acquired resistance (SAR) and that it plays an important role in plant defense against pathogens, including plant viruses (Vlot *et al.* 2009). Micropropagated peach shoots, in spite of high virus content, did not show any PPV infection symptoms (Clemente-Moreno *et al.*, 2011). Thus, we analyzed the effect of both MD and Phe treatments (using non-labelled compounds) on PPV-infected peach seedlings under greenhouse conditions. The presence of sharka symptoms in peach leaves was scored for each plant according to a scale of 0 (no symptoms) to 5 (maximum symptom intensity), a common test used in the evaluation of resistance to sharka (Decroocq *et al.* 2005; Rubio *et al.* 2005). According to the mean intensity of symptoms in peach leaves, we observed that MD-treated seedlings showed a significant decrease in PPV-induced symptoms (Fig. 6). This effect correlated with the increased SA levels found in MD-treated seedlings (Table 1). In contrast, although the Phe treatments also reduced sharka symptoms, no significant differences were observed compared with infected control seedlings (Fig. 6).

**Figure 6.**
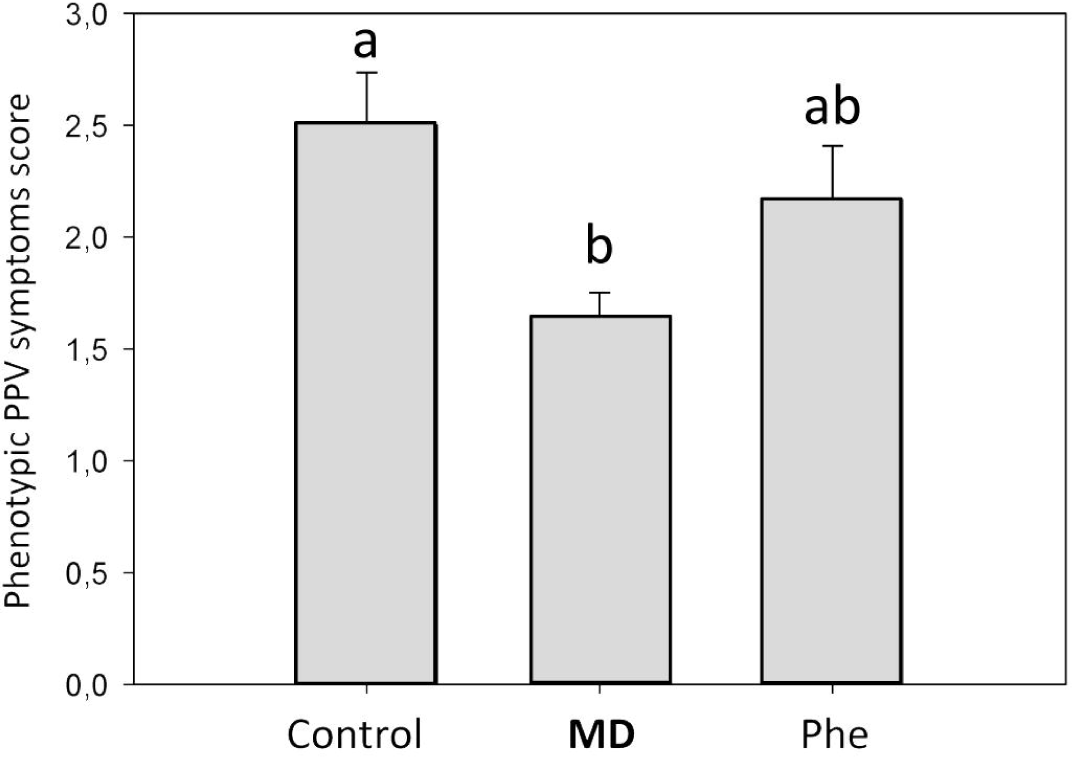
Phenotypic scoring for evaluating the resistance/susceptibility to PPV infection (Decroocq *et al*., 2005) and sharka symptoms in peach plants. Data represent the mean ± SE of at least 18 repetitions (samples from two independent assays carried out in 2015 and 2016) of each treatment. Different letters indicate significant differences according to Duncan’s test (P<0.05).

### Effects on antioxidative metabolism: the inhibition of H2O2-scavenging enzymes in MD-treated seedlings

The effects of both the MD and Phe treatments on the antioxidative metabolism of *in vitro* peach GF305 micropropagated shoots were analyzed. ANOVA analysis showed that the treatments had a significant effect on all the analyzed antioxidant enzymes except for peroxidase (POX) and superoxide dismutase (SOD). Micropropagated peach shoots treated with Phe displayed more significant increases in the enzymatic activities measured than control plants. MD treatments, on the other hand, significantly increased monodehydroascorbate reductase (MDHAR), dehydroascorbate reductase (DHAR), catalase (CAT) and SOD activities (Table 2). It is interesting to remark that Phe-treated plants seem to have more active ascorbate-glutathione (ASC-GSH) cycle enzymes than MD-treated micropropagated shoots.

**Table 2.** Effect of MD and Phe treatments on APX, MDHAR, DHAR, GR, POX, CAT, and SOD activities in *in vitro* GF305 shoots and in leaves of GF305 seedlings. APX, MDHAR, DHAR, and GR are expressed as nmol min^-1^ mg^-1^ protein. POX and CAT are expressed as |imol min^-1^ mg^-1^ protein, and SOD as U mg^-1^ protein. Data represent the mean ± SE of at least five repetitions. Different letters in the same column indicate significant differences according to Duncan’s test (P≤0.05). F-values from two-way ANOVA significant at the 99.9% (***), 99% (**), or 95% (*) level of probability (Treatment factor: Tto).

| *In vitro*<br>Treatment | APX | MDHAR | DHAR | GR | POX | CAT | SOD |
| --- | --- | --- | --- | --- | --- | --- | --- |
| Control | 316.0 ± 48.0 b | 866.2 ± 8.9 b | 246.0 ± 18.5 c | 342.8 ± 7.9 b | 1453.8 ± 22.4 b | 4.1 ± 0.1 b | 36.2 ± 2.5 b |
| MD | 257.0 ± 32.7 b | 1539.6 ± 81 a | 490.2 ± 29.6 b | 387.0 ± 8.7 b | 1600.4 ± 231.0 ab | 7.2 ± 1.1 a | 57.8 ± 8.1 a |
| Phe | 490.9 ± 35.9 a | 1538.6 ± 37.4 a | 791.3 ± 1.2 a | 489.6 ± 19.1 a | 1935.9 ± 78.6 a | 6.5 ± 0.4 a | 56.0 ± 3.4 a |
| ANOVA | F-values |  |  |  |  |  |  |
| Tto | 8.8* | 56.3*** | 190.7*** | 33.6** | 3.8 | 5.7* | 4.9 |

| *Seedlings*<br>Treatment | APX | MDHAR | DHAR | GR | POX | CAT | SOD |
| --- | --- | --- | --- | --- | --- | --- | --- |
| Control | 344.9 ± 64.6 a | 1262.9 ± 115.4 a | 472.8 ± 51.6 a | 180.4 ± 4.4 b | 1291.8 ± 136.9 a | 96.1 ± 6.8 a | 119.1 ± 17.1 b |
| MD | 198.2 ± 15.7b | 1300.7 ± 13.6 a | 405.7 ± 31.6 a | 196.3 ± 16.0 b | 969.1 ± 20.3 b | 53.25 ± 2.2 c | 107.9 ± 4.9 b |
| Phe | 403.6 ± 22.1a | 1106.6 ± 28.3 a | 436.2 ± 18.2 a | 283.8 ± 12.4 a | 1323.7 ± 53.0 a | 70.4 ± 1.9 b | 198.3 ± 12.9 a |
| ANOVA | F-values |  |  |  |  |  |  |
| Tto | 9.2** | 2.8 | 0.9 | 21.6** | 5.8* | 25.3** | 17.9** |

A different pattern was produced in GF305 seedlings. In this experiment, ANOVA analysis indicated that the treatments had a significant effect on ascorbate peroxidase (APX), glutathione reductase (GR), POX, CAT and SOD, but no significant effects were observed in the ascorbic acid-recycling enzymes (MDHAR and DHAR) in any of the treatments (Table 2). The MD treatment produced a significant decrease in APX, POX and CAT activities (H2O2-detoxifying enzymes), which could be associated with the observed increase in SA (Durner and Klessig, 1995; Rao *et al.*, 1997). The Phe treatment, on the other hand, increased GR and SOD activity and decreased CAT activity (Table 2).

We also studied the effect of MD and Phe on the ascorbate and glutathione content in leaf samples from peach seedlings. No oxidized ascorbate (DHA) was detected under our experimental conditions, and only the reduced form (ASC) was measured (Table 3). We observed that the treatments had a significant effect on ASC and oxidized glutathione (GSSG) concentrations as well as on the redox state of glutathione. Both treatments, particularly MD, decreased the ASC levels. In the case of MD, the decrease in ASC was about 50%, whereas Phe decreased ASC levels by about 30%. The effect of Phe or MD on reduced glutathione (GSH) was less pronounced: a 20% decrease was observed in both treatments (Table 3). In parallel, a significant accumulation of the oxidized form of glutathione (GSSG) was produced, mainly in Phe-treated plants, leading to a decrease in the redox state of glutathione in both MD- and Phe-treated plants (Table 3).

**Table 3.**
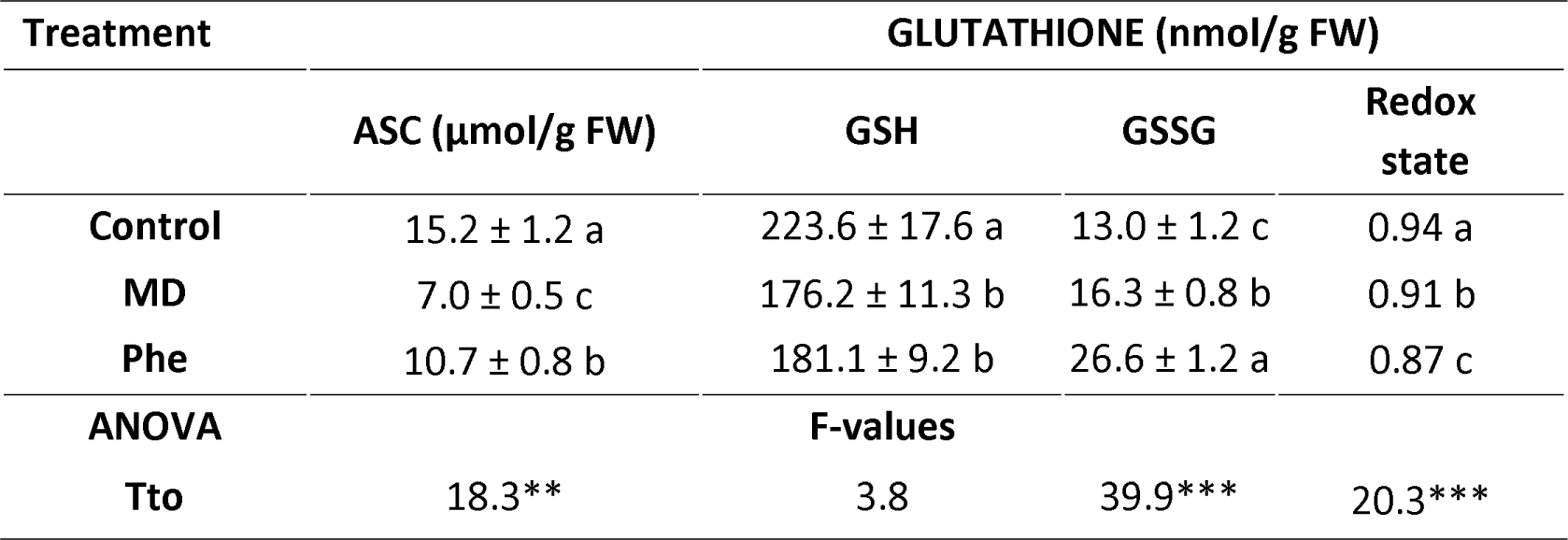
Effect of MD and Phe treatments on reduced ascorbic acid (ASC) and glutathione content in the leaves of GF305 peach seedlings. Data represent the mean ± SE of at least five repetitions. Different letters in the same column indicate significant differences according to Duncan’s test (P<0.05). F-values from two-way ANOVA significant at the 99.9% (***), 99% (**), or 95% (*) level of probability (Treatment factor: Tto).

### MDL1 activity and gene expression

Once we determined that the MD treatment increased MD, BA and SA in micropropagated peach shoots and SA levels in peach seedlings, we wanted to check if the MD could stimulate and/or up-regulate the mandelonitrile lyase (MDL) activity and/or the *MDL1* gene expression. MDL activity catalyzed the breakdown of MD into benzaldehyde plus cyanide (Swain and Poulton, 1994b). Benzaldehyde could then be oxidized by aldehyde oxidase into BA, an SA precursor. In micropropagated peach shoots, both the MD and Phe treatments significantly increased MDL activity (Fig. 4). Regarding *MDL1* gene expression, although a slight increase in gene expression was observed in MD- and Phe-treated micropropagated shoots, differences were not statistically significant (data not shown).

In peach seedlings, control and MD-treated plants had higher MDL activity levels than Phe-treated plants. Moreover, MD-treated peach seedlings showed a slight increase in MDL activity when compared with control plants, although differences were not statistically significant (Fig. 7A). Nevertheless, MD-treated seedlings displayed a 3-fold increase in *MDL1* gene expression (Fig. 7B).

**Figure 7.**
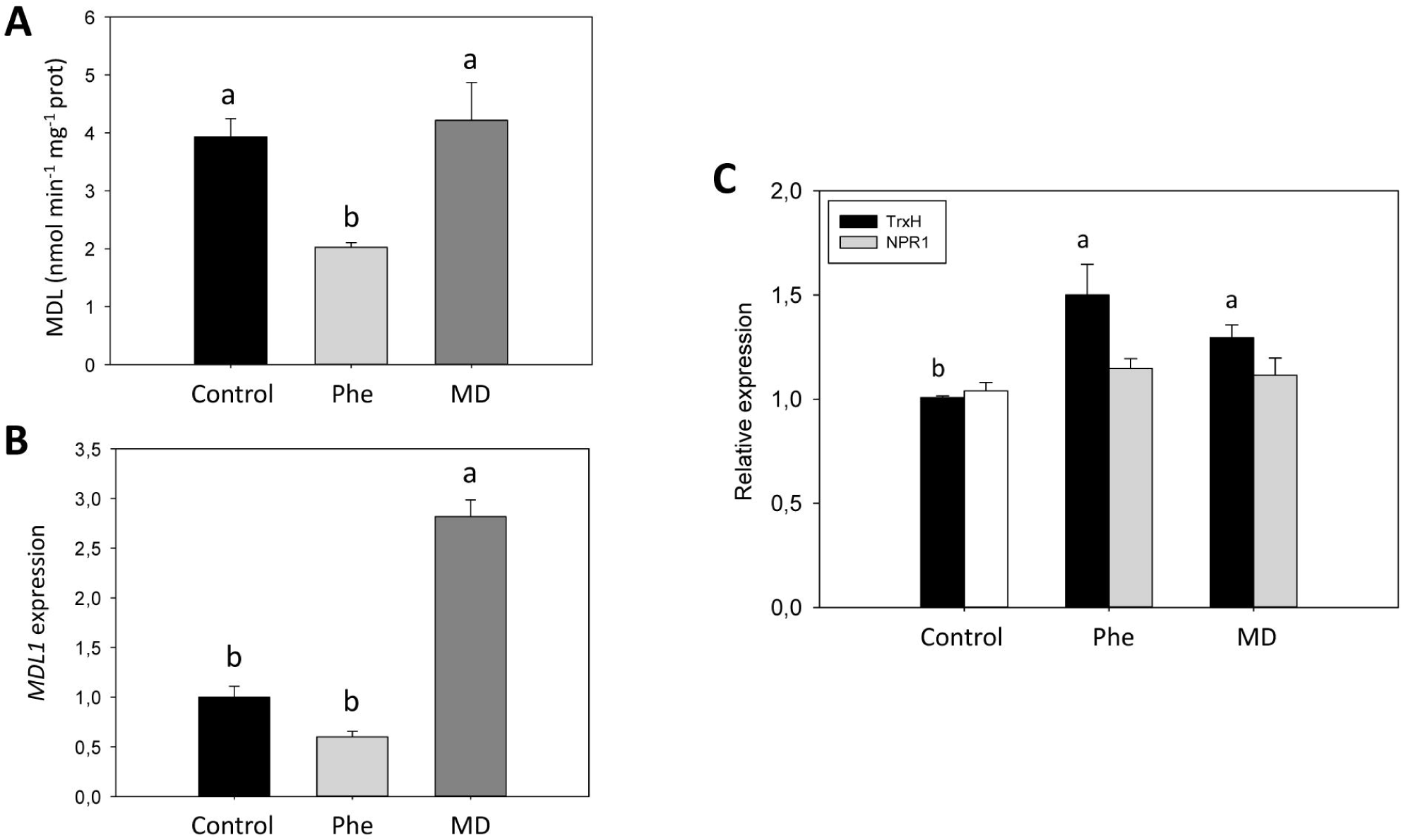
Mandelonitrile lyase enzymatic activity (A), *MDL1* gene expression (B), and gene expression of *TrxH* and *NPR1* (C) in GF305 peach seedlings grown in the presence or absence of MD or Phe. Data represent the mean ± SE of at least five repetitions of each treatment. Different letters indicate significant differences in each graph according to Duncan’s test (P≤0.05).

### Gene expression of redox-related genes

Due to the described role of SA in the induction of Non-Expressor of Pathogenesis-Related Gene 1 (*NPR1*) expression and the role of thioredoxins (Trx) in the SA-induced NPR1 conformational changes (Dong, 2004; Tada *et al.*, 2008; Vieira Dos Santos and Rey, 2006), we also analyzed the effect of both treatments (MD and Phe) on the *NPR1* and *TrxH* expression levels in peach seedlings. Whereas no significant changes in *NPR1* expression were observed with either treatment (Fig 7C), *TrxH* expression was significantly induced by Phe and MD treatments, with increases of about 42% and 23%, respectively (Fig 7C).

## Discussion

In this work, we have accumulated strong evidence suggesting that mandelonitrile could be metabolized into SA, linking the SA biosynthetic and CNglcs pathways in peach. Among this evidence, we observed increased levels of benzoic acid (BA) and SA as well as enhanced MDL activity and *MDL1* gene expression in MD-treated peach plants. Our results suggest that part of the total SA content in peach plants could be due to mandelonitrile, and hence CNglcs turnover. This possibility has not been described before in higher plants. Mandelonitrile seems to act as a hub in this pathway controlling both the amygdalin and SA biosynthesis. Similarly, in other plant species, the biosynthesis of the plant hormone auxin from tryptophan via cytochrome P450 enzymes has been also described (Mano and Nemoto, 2012).

The experiments with [^13^C]-labelled MD or Phe carried out using *in vitro* micropropagated shoots revealed significant incorporation of ^13^C as [^13^C]SA and [^13^C]BA, one of the SA precursors, in [^13^C]MD-treated micropropagated shoots. In fact, almost 40% of total BA and 20% of total SA appeared as ^13^C-labelled compounds. In addition, and as expected, an important increase in amygdalin content was observed in both treatments. However, a possible route from Phe to BA cannot be ruled out because a slight increase in BA was observed in [^13^ C]Phe-treated micropropagated shoots, although this possibility does not seem to be important for SA biosynthesis because no parallel increase in [^13^C]BA or in SA was recorded. On the other hand, Phe is a ketogenic and glucogenic aminoacid, and it is a precursor for a wide range of specialized natural compounds (Yoo *et al.*, 2013), including CNglcs, in peach plants. Moreover, treatment with [^13^C]Phe increased the percentage of [^13^C]MD observed, highlighting the central role MD plays in controlling both SA biosynthesis and CNglcs synthesis and turnover.

It is known that during SA biosynthesis, benzaldehyde can be oxidized into BA in a reaction catalyzed by aldehyde oxidase (Dempsey *et al.*, 2011). In addition, the results for MDL activity and *MDL1* gene expression reinforce the possible role of MD in the increase in SA via BA. Accordingly, we observed a high *MDL1* expression level to sustain increased MDL activity when peach seedlings were treated with MD, but not in Phe-treated plants. We do not know the exact cellular location in which these reactions take place. In *Prunus serotina*, MDL was immunocytochemically localized in cell walls and vacuoles (Swain and Poulton, 1994a). However, no studies of the sub-cellular localization of MDL have been conducted in peach leaves. In previous works, due to the low contamination levels obtained, we suggested an apoplastic localization of MDL (Diaz-Vivancos *et al.*, 2006), although a vacuolar localization should not be excluded.

Using a different analytical method than that used for micropropagated shoots, we were also able to detect an increase in leaf SA content under MD treatment in peach seedlings. Curiously, the SA increase observed was similar in both plant models (88% and 83% in peach seedlings and micropropagated shoots, respectively). The MD treatment, besides increasing SA, also enhanced the concentration of ABA, GA1 and DHZ. It is known that cross-talk among different hormonal signals is involved in different physiological responses as well as in response to environmental challenges (Grant and Jones, 2009; Micol-Ponce *et al.*, 2015; Yang *et al.*, 2013). Among plant hormones, ABA and SA, along with JA and ethylene, play a major role in mediating plant defenses against biotic and abiotic factors (Verma *et al.*, 2016; Yang *et al.*, 2013). Moreover, the interaction of SA with other plant hormones regulating certain plant responses has been also reported (Bari and Jones, 2009). For example, the role of cytokinins in modulating SA signaling in biotic and abiotic stress responses is well documented (Choi *et al.*, 2010; Rivero *et al.*, 2009). GAs levels and signaling are also involved in plant defense (Zhu *et al.*, 2005), and the interaction of auxins with SA can limit disease through the down-regulation of auxin signaling (Wang *et al.*, 2007). ABA can also promote plant defense responses depending on different factors, such as the pathogen, the development stage of the plant and the target tissue (Yang *et al.*, 2013).

NPR1 is a key regulator in the signal transduction pathway that leads to SAR. The induction or overexpression of the NPR1 protein leads to increased induction of *PR* genes and enhanced disease resistance (Dong, 2004). Inactive NPR1 occurs in the cytosol as oligomers, held together by disulfide bridges. SA-induced changes in the redox state can lead to monomerization by the reduction of Cys residues via the action of TrxH (Tada *et al.*, 2008) (Fig. 7). NPR1 monomers are translocated to the nucleus, activating defense genes (Vlot *et al.*, 2009). Under our experimental conditions we were not able to detect any *NPR1* induction, although the up-regulation of *TrxH* was noticed, suggesting its role in the activation of NPR1 monomerization and thus in enabling the activation of defense genes. Similarly, in a previous work, we were not able to detect any *NPR1* gene induction in micropropagated peach shoots treated with benzothiadiazole, a SA analogous inducer of SAR (Clemente-Moreno *et al.*, 2012). The interaction of SA with heme-containing proteins, such as the H2O2-scavenger enzymes (CAT, APX or POX), can produce redox stress, which can initiate the release of NPR1 monomers and their entry into the nuclei (Durner and Klessig, 1995). Accordingly, we described a decrease in these enzymes in MD-treated peach plants. Moreover, MD also decreased the levels of reduced ascorbic acid and the glutathione redox state, leading to a more oxidized environment (Fig. 8). It has been suggested that SA induces changes to the redox environment by modulating GSH levels and reducing power, stimulating the plant defense responses (Herrera-Vasquez *et al.*, 2015; Vlot *et al.*, 2009; Yang *et al.*, 2004). The known role of SA and ABA in the control of stomatal closure (Khokon *et al.*, 2011) and the MD-induced SA and ABA levels (Table 1) displayed in peach seedlings could led us to speculate that MD-treated peach plants could tolerate situations of water and/or saline stress.

**Figure 8.**
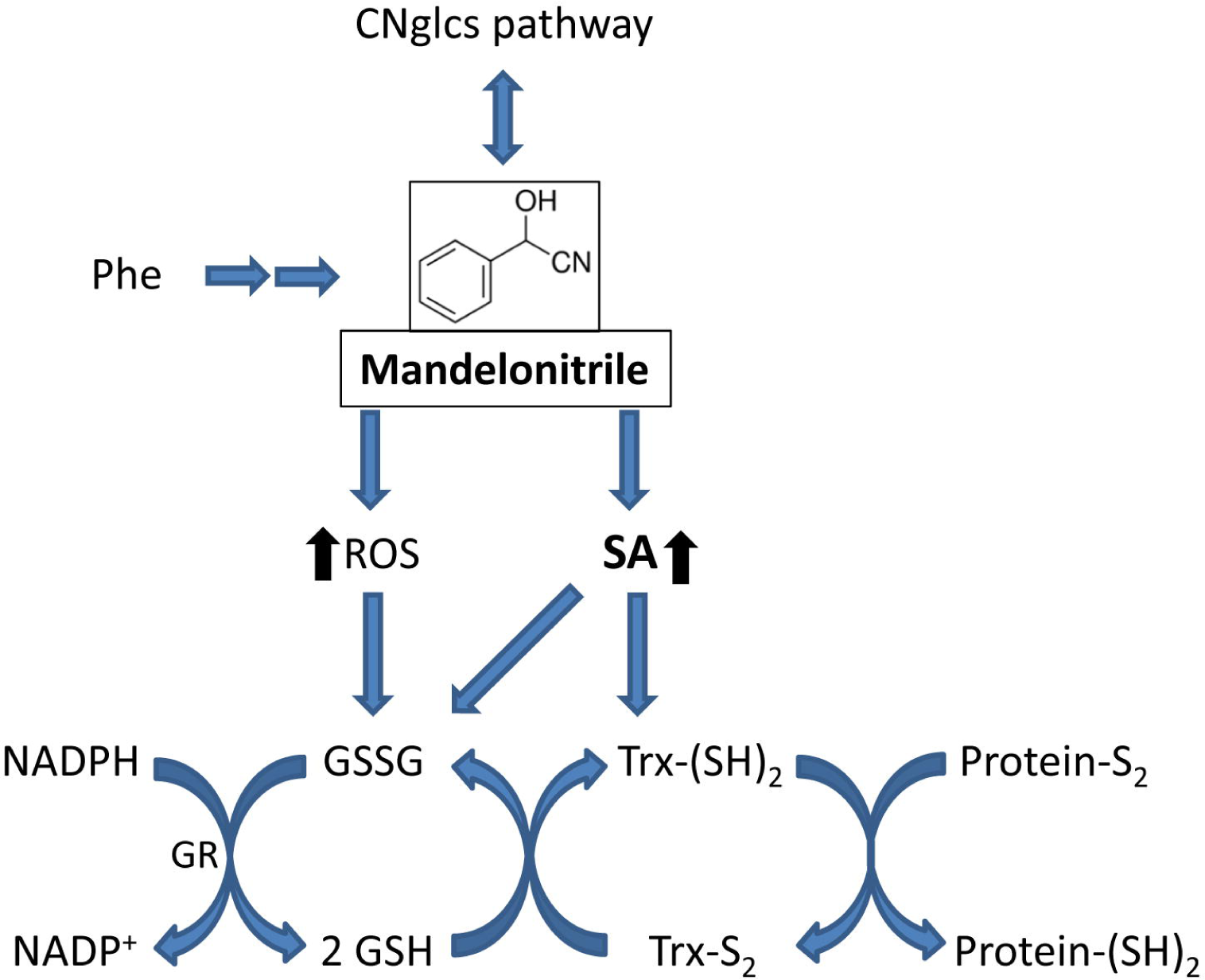
Proposed roles of MD in peach plants. MD is involved in CNglcs turnover and in SA biosynthesis. In addition, MD treatment leads to a more oxidized environment, which could modify the function of proteins such as those involved in the response to environmental stress conditions.

We have observed that the MD treatment induced partial protection against PPV infection in peach seedlings. Based on our results, we can suggest that the partial protection from PPV in MD-treated GF305 peach plants could be independent of *NPR1* induction. Indeed, it is known that some SA-induced defense genes do not require *NPR1*, suggesting that other proteins can be important in SA perception (Blanco *et al.*, 2005) (Fig. 8). Moreover, the MD treatment increased DHZ levels, and this cytokinin has been found to induce partial protection against *White clover mosaic virus* with no changes in the expression of SA-responsive genes (Gális *et al.*, 2004). Moreover, in spite of its well-known role on plant defense against pathogens (Alvarez *et al.*, 1998), SA has been increasingly recognized as abiotic stress modulator via SA-mediated regulation of important plant-metabolic processes (Khan *et al.*, 2015). Thus, MD levels and hence the CNglcs turnover could be involved in defense responses by increasing the levels of SA and/or the interaction with oxidative signaling defense-induced pathways. On the other hand, in micropropagated peach shoots, although MD treatment did not increase the SA content under abiotic and biotic stress conditions (Fig. 5), data from Fig. 3 show that a small amount of MD is metabolized to SA. Taking together, we suggest that under stress conditions this new SA biosynthetic pathway contributes much less to the total amount of SA than the PAL pathway.

We have therefore shown that the addition of a small molecule like MD can reveal very useful information on the mechanisms of interaction of various hormones in coordinating responses to environmental stress in plants. In conclusion, in this work we provide strong evidence for a new SA biosynthetic pathway from MD in peach by using two different plant models and different analytical approaches. Although we acknowledge that additional genetic evidences would provide complementary data, the feasibility of genetic approaches in peach plants is scarce nowadays. The MD molecule seems to act as a hub in this novel pathway controlling amygdalin and SA biosynthesis (Fig. 8). The MD treatment induced the gene expression of *MDL1* to maintain MDL activity. This result reinforces the possible role of MD in the increase of SA via BA, which has been described as an SA precursor (Dempsey *et al.*, 2011; Ribnicky *et al.*, 1998). In addition, there was also a pleiotropic effect on other plant hormones (ABA, GA1, DHZ). MD, and therefore SA, induced *TrxH*, but not *NPR1*. However, the effect of MD (or SA) in the antioxidative machinery can induce redox stress, which can facilitate NPR1 monomerization and therefore its effect on defense gene induction (Durner and Klessig, 1995). This argument is supported by the partial protection against PPV induced by the MD treatment. Alternatively, and based in our results, it is also possible that the partial protection against PPV in MD-treated GF305 peach plants could be independent of *NPR1* induction, as described by other authors (Blanco *et al.*, 2005). Nevertheless, because this new pathway seems not to be relevant under stress conditions, at least under *in vitro* conditions, further investigation will be required in order to elucidate how relevant is this new pathway for plant performance.

## Supplementary data

**Fig. S1.-** Effect of [^13^C]MD or [^13^C]Phe on NaCl-stressed micropropagated peach shoots.

## Acknowledgments

This work was supported by the Spanish Ministry of Economy and Competitiveness (Project AGL2014-52563-R). PDV and CP thank CSIC and UPCT, respectively, as well as the Spanish Ministry of Economy and Competitiveness for their ‘Ramon & Cajal’ research contract, co-financed by FEDER funds. We also acknowledge Prof. Manuel Acosta Echeverría for his very useful commentaries and discussion.

